# Quantitative evaluation of LED based optical autofocus module

**DOI:** 10.64898/2026.04.10.717415

**Authors:** S. Habte, S. Kumar, J. Lightley, E. Garcia, M.A.A. Neil, P.M.W French

**Affiliations:** Physics Department, Imperial College London, London SW7 2AZ, UK; Francis Crick Institute, 1 Midland Road, London, NW1 1AT, UK

## Abstract

We report an improved version of the open-source optical autofocus module (“*openAF*”) for light microscopy using a light emitting diode (LED), together with a method to independently quantify the performance of optical autofocus systems using 2D autocorrelation analysis of astigmatic imaging of fluorescent nanobeads. We apply the latter for both the LED-based and the previous super luminescent diode (SLD) based implementations of the *openAF* optical autofocus approach used in conjunction with a 100x 1.4 NA oil-immersion objective lens. The new approach accounts for power variations in the autofocus light source and we demonstrate that the convenient LED-based system can provide axial stability with a standard deviation <10 nm over at least 45 minutes when switched on from cold, during which the LED power varies as it reaches thermal equilibrium.

## Introduction

Autofocus (AF) techniques are essential in optical microscopy to ensure that the sample remains in focus despite mechanical or environmental perturbations such as thermal drift, and/or variations in distance from objective lens to coverslip. AF is a basic requirement of automated microscopy, e.g., for high-content analysis (HCA) and whole-slide scanning – noting that throughput is an increasing priority for light microscopy, e.g., for automated pathology and the increasing requirements for large experimental data sets for training machine learning models, e.g., ^1,2^.

As well as high throughput or long timescale imaging at high numerical aperture (NA), such as time-lapse assays, AF is also important for super-resolution microscopy. Single molecule localisation microscopy (SMLM)^3,4,5^ can achieve nanometre-scale resolution but requires data acquisitions of many minutes, during which the sample can drift in three dimensions. Although lateral drift can be corrected post-acquisition via spatial cross-correlation of image data (including with the use of fiduciary markers)^6,7^, axial drift by more than the depth of field can cause point spread function (PSF) blurring and thus localisation errors that degrade reconstructed super-resolved images. For 3D SMLM, which may be implemented, e.g. using methods such as astigmatism^8^, biplane^9^ and engineered PSFs^10^ that rely on axial distance measurement via the calibrated variation of the PSF, the axial resolution can also be compromised by drift in focus.

AF can be implemented utilising existing motorised microscope functionality with software-based approaches, or by the use of additional optical hardware modules for optical AF (OAF). Software-based AF techniques typically rely on image quality metrics of the sample image - such as sharpness, entropy or spatial frequency content^11,12,13^ - that are evaluated across a reference axial image stack for correlation with the test image metric to determine defocus. However, this reference z-stack typically needs to be acquired for each different sample or field of view (FOV) - significantly slowing image data acquisition rate for slide scanning or multiwell plate assays. Moreover, such image metric-based approaches work best with thin samples and may fail with extended samples such as tissue slices or 3D cell cultures, and for samples that evolve over time.

OAF approaches utilise an optical beam (often in the infrared spectral region to minimise both phototoxicity and overlap with fluorescence signals) that is reflected from one of the microscope sample-coverslip interfaces during imaging. Defocus can be quantified via position or image-based metrics of the reflected AF light, typically detected using a photodiode array or a camera, such as the autofocus beam’s intensity, displacement or size^14,15,16^, or the autofocus camera image file size^17^. These approaches also entail the acquisition of a “calibration” z-stack AF data set to which the test value of the defocus metric can be compared. However, this is usually required only once in an imaging session (for a specific microscope configuration) and generally does not need to be repeated for each different sample or FOV. Informed by the calculated defocus, the sample or objective lens can be continuously moved to bring the sample back to focus. OAF can provide faster operation, e.g., for real-time “closed loop” operation, and extended axial range compared to image-based methods and can operate with reduced phototoxicity since the AF is independent of the primary imaging process and may utilise low power infrared radiation.

A hybrid AF approach is to use features in the sample^18^ or fiduciary markers ^19,20^ in the real time images, such as sub-resolution polystyrene beads or gold nanorods affixed to the coverslip, to enable simultaneous lateral and axial position locking or 3D drift-correction using either the primary imaging detector or a separate AF imaging system. However, this approach requires additional sample preparation and/or complex computational analysis to realise 3D drift-correction, which may include fitting, e.g., ^21^, or deep learning, e.g., ^22^, to account for field dependent optical aberrations.

Although several OAF configurations have been reported, e.g., reviewed in ^23^, including AF capabilities built into commercial microscope options^24,25^, there is a paucity of robust AF modules available for after-market implementation on custom-built or commercial microscopes. With the growing number of custom microscopes being developed for diverse applications, there is a need for robust, adaptable and cost-effective OAF modules, including open-source approaches, that offer straightforward implementation and flexibility. Thus, the development of robust OAF implementations remains an active field of research, noting that even integrated commercial AF systems can fail with challenging samples or specialised experimental requirements. One consideration is that thermal or mechanical drifts in microscopes in the AF systems themselves can also impact the microscope performance without being detected or corrected. A further challenge is that, since the intensity of the back-reflected AF signal depends on the Fresnel reflection at the coverslip interface, OAF can be challenging to implement when imaging biological samples with high-NA oil immersion objective lenses since the reflectivity from the coverslip/aqueous medium interface is only ∼0.4% and this signal may be weaker than unwanted back-reflections from other surfaces or light backscattered from the biological sample itself. Thus, a practical AF implementation needs to be robust against drift in the autofocus system itself, and against unwanted back-reflections and other spurious signals incident on the AF detector.

To develop a robust OAF, it is important to quantity the performance independently of the OAF system, preferably with a rapid and convenient method. In previous work from our laboratory^15,26^, we analysed z-stacks of fluorescent beads (using the software *PSFj*^27^)) to determine the net defocus after operation of our OAF, and also used the sharpness of a transmitted light image of a USAF test chart as a readout of defocus. However, the repeated acquisition of bead stacks to provide a defocus metric was time-consuming, and the use of a USAF test chart provided an anomalously strong back reflection of the AF beam that may not recapitulate the performance experienced when imaging biological samples. More recently an astigmatism-based OAF utilising a laser diode, *PiFocus*^*16*^ was reported that used the astigmatic 3D localisation method of a bead sample to characterise time-lapse performance of their OAF – demonstrating <10 nm precision.

In this paper, we review our development of OAF modules for the modular, sustainable and open-source microscope platform, *openScopes*^*28*^ that we have designed to enable rapid prototyping by instrument developers and to widen access to research-grade microscopy for users in lower resourced settings. These open-source OAF implementations are intended to be applicable to most fluorescence microscopes, including for SMLM, HCA and slide-scanning, and are specifically designed to work with *openFrame*-based instruments^26^, which can support advanced imaging modalities including automated SMLM based on *easySTORM*^29,15,^ and quantitative phase imaging using single-shot *pDPC*^30,31,^.

Our first implementation of an OAF^15^ utilised a single mode laser diode collimated by a spherical lens and a rectangular slit was used to provide an autofocus beam with different orthogonal confocal parameters when focussed on the microscope coverslip. The back reflected beam was then focussed onto a CMOS camera with the aim of using the beam size to provide the defocus metric. We reasoned that this would be more robust against drifts in alignment that could impact metrics based on autofocus beam position. In principle, this approach enables the OAF to provide higher resolution when considering the beam projection parallel to the slit (since the beam diameter at pupil of objective lens is larger), and longer range when considering the perpendicular beam projection. Because the axially varying AF camera images were compromised by interference between the AF signal and unwanted back-reflections, we used a convolutional neural network (CNN) trained on calibration AF camera z-stacks acquired over multiple days to determine the defocus from the camera image data, rather than an analytic algorithm^15^. This also yielded the sign of defocus from a single test image. This CNN-based OAF approach achieved ±100 µm operation range with an accuracy better than the depth of field (DOF) of the 1.4 NA objective lens using a two-step algorithm. However, significant time was required to obtain the composite CNN training data over ∼10 days (necessary to account for routine perturbations in the alignment of the AF system itself) and the training data z-stacks had to be sampled to the desired precision of the OAF, which entailed large data volumes. While this CNN-based OAF worked reliably for automated (multi-well plate) SMLM assays over >12 months, the requirement to acquire new training data for each specific microscope configuration was a practical disadvantage.

Accordingly, we developed an analytic approach analysing the AF camera image data to realise “*openAF*”^26^ where we replaced the combination of spherical collimation and a slit with collimation of the light from a single mode optical fibre using two orthogonal cylindrical lenses of different focal lengths. One of these cylindrical lenses is slightly translated from the collimation position to provide a shift in the focal plane at the autofocus camera. This enables the sign of defocus to be determined from a single test image. We also replaced the AF laser diode with a super luminescent diode (SLD) that greatly reduced the interference of the AF signal with spurious reflections such that the incoherent background did not vary axially (i.e., with defocus) and could be subtracted from each AF camera image. The defocus was determined by calculating the width of the Fourier transform of the average image intensity projection of the recorded autofocus camera image along each orthogonal axis. This *openAF* module was implemented on an *openFrame*-based (*easySTORM*) microscope^26^ to realise single-shot closed loop operation with an axial range of ± 37 µm with <200 nm accuracy (measured using bead z-stacks) and could be operated in a two-step autofocus mode with an axial range up to ± 68 µm. While the *openAF* module provides a robust OAF capability that has been used for SMLM, including for super-resolved CLEM^32^, the SLD is a relatively expensive and fragile component, and both SLDs and laser diodes present a laser safety issue that must be carefully managed.

Here we report further development of the *openAF* module where the single mode optical fibre (SMOF)-coupled SLD has been directly replaced with a low-cost multimode optical fibre (MMOF)-coupled LED – with other elements of the *openAF* module remaining the same. As well as being easier and cheaper to acquire, the use of an LED light source removes the need to address laser safety in the installation and use of an OAF module. To quantify the performance of this new OAF, we implemented astigmatic fluorescence imaging on the *openFrame*-based microscope by inserting a cylindrical lens (CL) in the fluorescence imaging path, as is commonly used for 3D SMLM. To provide rapid and straightforward quantification of its performance we acquired time-lapse (astigmatic) image data of fluorescent nanobeads and developed a novel image autocorrelation-based software tool to compute the defocus independently of the OAF readout. Since this works with a single fluorescence bead image, it is much simpler to implement than acquiring bead z-stacks, and the computational analysis is much faster than 3D (SMLM software-based) localisation of individual beads. When characterising our LED-based openAF module, we observed an LED power-dependent artefact that was apparent as the LED reached thermal equilibrium after being switched on. We therefore modified the *openAF* algorithm to include a normalisation of the detected autofocus light intensity and subsequently achieved <10 nm precision from switch-on over extended time-lapse measurements (∼ 45 minutes).

During this work we also developed an “infinity alignment tool” that provides a simple means to optimise focusing of the imaging arm (tube lens to camera) of a microscope, including where astigmatic imaging has been implemented.

## Methods

### LED based *openAF*

Figure 1 presents a schematic of the modified *openAF* autofocus module coupled to the *openFrame*-based microscope^26^. Here the SLD has been replaced with an 850 nm LED (Thorlabs M850F1, with LEDD1B driver) coupled into a 50 µm core diameter multimode optical fibre (Thorlabs M14L). This extended optical source with low spatial coherence is not as well-collimated as the single spatial mode SLD beam coupled via a SMOF and the resulting image on the autofocus camera includes light emerging from the MMOF cladding as well as the core. There is also significant unwanted scattered light within the autofocus module and so we introduce a 3D printed 3×10 mm rectangular aperture in front of the collimating cylindrical lenses to block unwanted radiation emerging from the cladding. Supplementary video 1 shows how the autofocus camera image varies as the microscope coverslip is scanned through focus and Supplementary Figure 1(a) shows autofocus camera images acquired when the sample coverslip is at defocus values of -10, 0 and +10 µm. For comparison, Supplementary Figure 1(b) shows analogous autofocus camera images when the SLD/SMOF are used. The OAF calculates the microscope defocus from the spatial profiles of these images, comparing the derived autofocus image metrics to those for a reference calibration data set that can be acquired within a few minutes at the start of each imaging session. These calibration data are obtained by determining the defocus metric value for each background-subtracted autofocus camera image in a z-stack of images acquired at known stage displacements from the focal plane and interpolating them to produce smooth curves.

**Figure 1.**
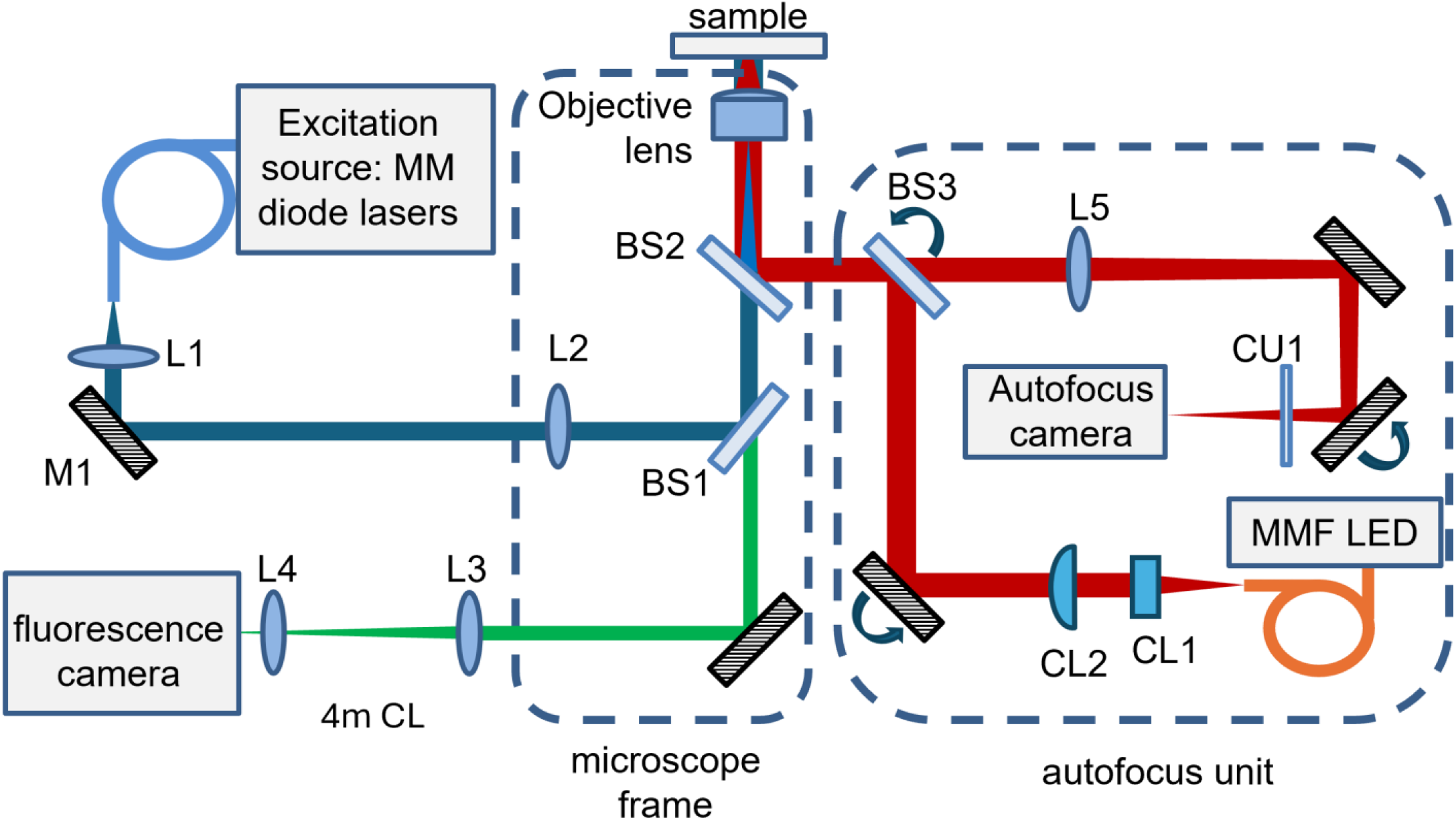
Schematic of openFrame-based microscope with LED-based optical autofocus: Excitation source – multimode diode laser bank (Triline, Cairn Research Ltd), L1 – 60 mm collimating lens (AC254-060-A, Thorlabs), L2 – 200 mm focusing lens (49-364, Edmund Optics), BS1 – multiline dichroic (ZT405_465_625_825rpc-UF2, Chroma), Objective lens – 100x 1.4 NA Oil objective lens (UPLSAPO) mounted on a piezoelectric actuator (Piezo Z-stage Q-545.140 with controller E-873.1AT, PI) to prove axial translation, L3 – 200 mm tube lens (MXA20696, Nikon), 4 m focal length cylindrical lens (4000 YO 65, Comar Optics, Ltd), L4 (0.35x Motic c-mount adapter (1101001904111, Motic), fluorescence camera (CellCam Kikker, Cairn Research Ltd). Autofocus unit: 850 nm LED (Thorlabs M850F1 with LEDD1B driver) coupled into 50 µm core diameter MMOF (Thorlabs M14L), CL1 – 6.4 mm focal length cylindrical lens (Thorlabs LJ1227L1-B, Thorlabs), CL2 – 20 mm focal length cylindrical lens (LJ1960L1-B, Thorlabs), L5 – 200 mm focal length lens in autofocus unit (AC254-200-B, Thorlabs), BS2 – 800 nm short pass autofocus dichroic (69-220 Edmund Optics), BS3 – 50:50 beamsplitter (47-026 Edmund Optics), CU1 – cleanup filter (FF01-842/56-25, Semrock), Autofocus camera (CM3-U3-31S4M-CS, FLIR).

Figure 2 shows the defocus calibration curves of three defocus metrics calculated from a background-subtracted autofocus camera z-stack acquired as the 100x 1.4 NA oil immersion objective lens (Olympus UPLSAPO) was translated through focus from -15 to +15 µm: the variation of the average autofocus camera image intensity (green) and the full width at half maximum (FWHM) of the Fourier transforms of the average image intensity projections of the recorded signal for each orthogonal axis (red and blue). These “calibration” autofocus z-stack image data were acquired in steps of 100 nm together with a background image recorded with the objective lens translated to 50 µm below focus. The calibration data acquisition took 2 minutes for this ±15 µm range. In operation, the *openAF µManager* plug-in autofocus module continuously acquires autofocus camera images, from which the background image is subtracted before computing the defocus metrics and comparing them to the values in the calibration look-up table. In closed loop operation the FWHM X (parallel to the axis of the longer focal length cylindrical lens CL2) metric is used to calculate the two possible defocus values (either side of the peak) through cubic spline interpolation of adjacent values, and the corresponding FWHM Y is used to determine the true defocus value. In the cases where the defocus would be near (at ∼±2 µm) the maxima of the FWHM X and FWHM Y curves, the orthogonal FWHM metric is used to determine defocus. The *µManager* autofocus plug-in^26^ then moves the objective lens to correct the calculated defocus.

**Figure 2.**
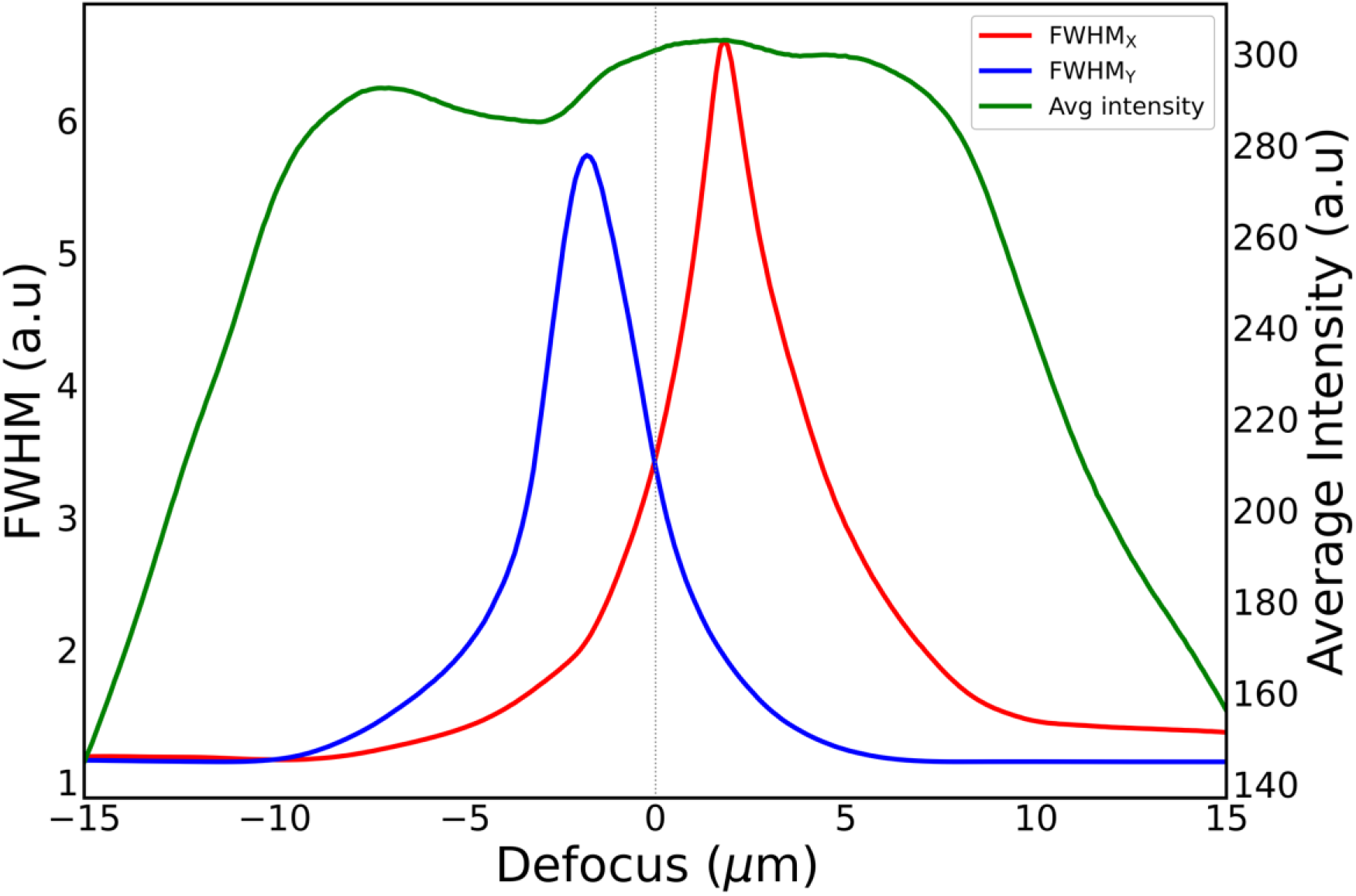
LED openAF axial calibration curves acquired between ±15 µm: interpolated plots of FWHM X (red) and FWHM Y (blue) of the of the Fourier transforms of the average image intensity projections along Y and X respectively, with the average intensity (green).

The average camera intensity decreases as the image of the LED radiation emerging from the MMOF becomes out of focus. Because the quasi-collimated LED/MMOF beam diverges much more rapidly than the collimated SLD/SMOF beam, the total operating range of the LED/MMOF autofocus is lower – at ∼±15 µm range. This is adequate for time-lapse imaging at a given field of view (FOV) and the autofocus can be operated in closed loop over this range. Typically, the autofocus camera is set to acquire images with an integration time of ∼100 ms and the total time to acquire each image, calculate the defocus and send the correction command to the objective lens z-stage is ∼250 ms.

### Measurement of autofocus performance over time using astigmatic imaging

To quantify the performance of this LED/MMOF-based *openAF* implementation and compare it with the previously published SLD/SMOF implementation, we measured the microscope’s defocus over time independently of the autofocus system. We acquired time-lapse images of 100 nm diameter fluorescent beads (TetraSpeck, T727) with a cylindrical lens of 4 m focal length inserted between the fluorescence imaging tube lens and the fluorescence camera assembly -which included a 0.3x Motic demagnifier mounted directly onto the (CellCam Kikker, Cairn Research Ltd) camera in order to permit imaging at a higher frame rate^26^. This astigmatic imaging would enable the fluorescent beads to be axially localised following the usual 3D SMLM approach^33^. However, since we are only interested in a single defocus value for the fluorescent bead image, rather than localisation of individual fluorescent beads, we computed the 2D autocorrelation function of the bead images across the FOV through the Wiener-Khinchin theorem, as illustrated in Supplementary Figure 2. We derived a defocus metric based on the ratio of the widths of Gaussian functions fitted to the orthogonal (“x” and “y”) mean intensity projections of the central peak of the 2D autocorrelation function (Supplementary Figure 2(c)). The protocol to acquire the calibration data for this astigmatic imaging defocus model is given in the Supplementary Information.

Supplementary videos 2 and 3 show the z-stack of acquired fluorescent bead images and of the corresponding maximum peak of the 2D autocorrelation function respectively. Supplementary Figure 2 shows exemplar plots of the maximum of the 2D autocorrelation functions at different values of defocus with the mean intensity projections along X and Y. Figure 3(a) shows the standard deviation, σ, of the Gaussian fits to the mean intensity projections along X (red) and Y (blue) of the central autocorrelation peak as a function of defocus, and Figure 3 (b); the ratio σ_X_/σ_Y_. The latter provides a monotonic range for precise interpolation and direct look-up table of defocus from the autocorrelation function of a given fluorescent image within the range of approximately ± 0.7 µm, defined between the extrema of the ratio plots. Thus, this analysis of astigmatic fluorescence images of a fluorescent bead sample provides a rapid means to independently quantify defocus of the microscope image and evaluate the performance of (any) autofocus module.

**Figure 3.**
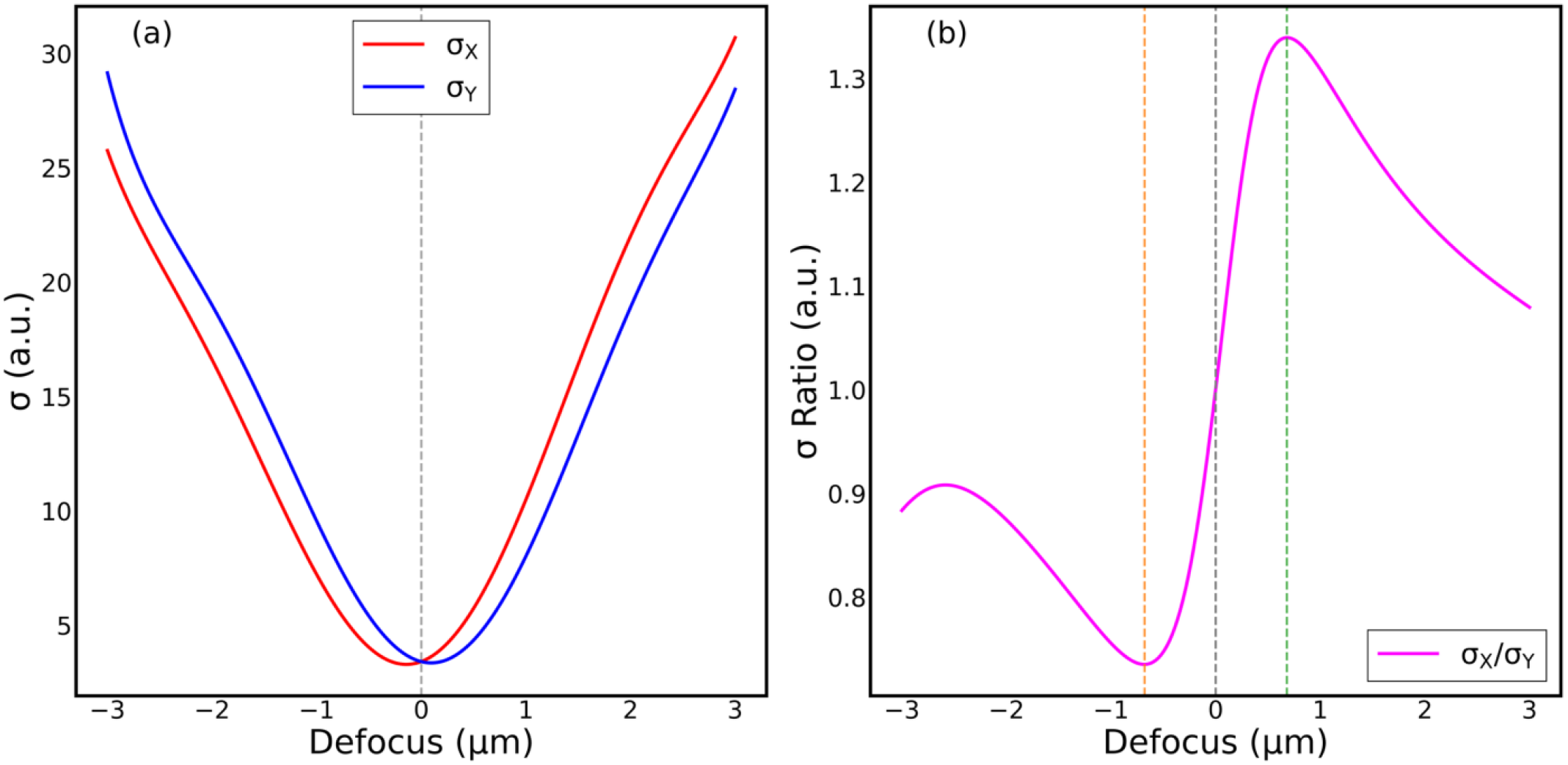
Astigmatic autocorrelation defocus model: (a) σ_X,Y_ of the 1D Gaussian fit to mean intensity projections in X (red) and Y (blue) of the central autocorrelation function peak; (b) Ratio σ_X_/σ_Y_ of Gaussian fits to Y and X mean projections of the central peak of 2D autocorrelation function where the interpolated monotonic range between the extrema (∼±0.7 µm range) is used for direct estimation of defocus value.

### Infinity alignment tool

We note that this characterisation of the performance of the OAF requires careful alignment and focusing of the fluorescence and autofocus imaging systems. To independently determine the conditions of good focus for each system, we developed the “infinity alignment tool” shown in Supplementary Figure 3. This comprises a metal tube with a graticule (R1DS3N2, Thorlabs) fixed at the focal plane of a 4x magnification objective lens RMS4x, Thorlabs) that could be screwed directly into the microscope objective holder (replacing the primary microscope objective lens) and so present an image at infinity to the tube lens. The fluorescence imaging camera can then be set to the position corresponding to the sharpest image of the graticule (illuminated with a convenient diffuse light source) and the primary microscope objective lens can then be replaced and translated to locate the (fluorescent bead) sample in the focal plane. The same infinity alignment tool can also be used to set up the separation of the autofocus tube lens L5 and the autofocus camera.

To precisely set the graticule in focal plane of the 4x objective lens in the infinity alignment tool, we use a shearing interferometer (SI100, Thorlabs) to check the collimation of an alignment laser beam that is retroreflected from the graticule surface after passing through the 4x objective lens – a procedure that is only required once. We believe this alignment tool is generally useful to rapidly set up any light microscope. For astigmatic imaging systems, such as we implement here and which are often used in 3D SMLM microscopes, the cross at the centre of the graticule can be used to determine the camera position that brings each axis of the astigmatic imaging system into focus – and then the camera can be set equidistant between them, at the circle of least confusion – as can be confirmed by observing equal defocussing of the graticule ruling in each orthogonal direction.

## Results

The astigmatic fluorescent bead imaging method was first used to monitor the accuracy of the LED *openAF* autofocus module where the LED was coupled into a 50 µm core diameter MMOF and the sample of 100 nm diameter fluorescent beads was imaged with 100x 1.4 NA il immersion objective lens (UPLSAPO). We observed that, while the autofocus generally functioned well, there could be a significant drift in focus after the LED was switched on. This behaviour is illustrated in Figure 4, which presents the time-lapse plots of (a) the defocus measured with the astigmatic bead imaging (magenta), (b) the defocus correction applied by the LED/MMOF *openAF* module (red), (c) the variation in power of the LED as represented by the average intensity at the autofocus camera (green) and (d) the position of the piezo-electric actuated z-stage reported by its encoder (purple). It is apparent from (c) that the LED power decreased by ∼2% during the ∼20-minute warm-up period after switching on and this led to a drift in the image focal plane of >500 nm before exceeding the range of the astigmatic imaging after 20 minutes. However, this focus drift caused by the LED power change is not detected by the *openAF* as reported by the required defocus correction (b). This figure illustrates the importance of independently measuring the imaging defocus to quantify the performance of an optical autofocus system during operation, rather than relying solely on the optical autofocus readout.

**Figure 4.**
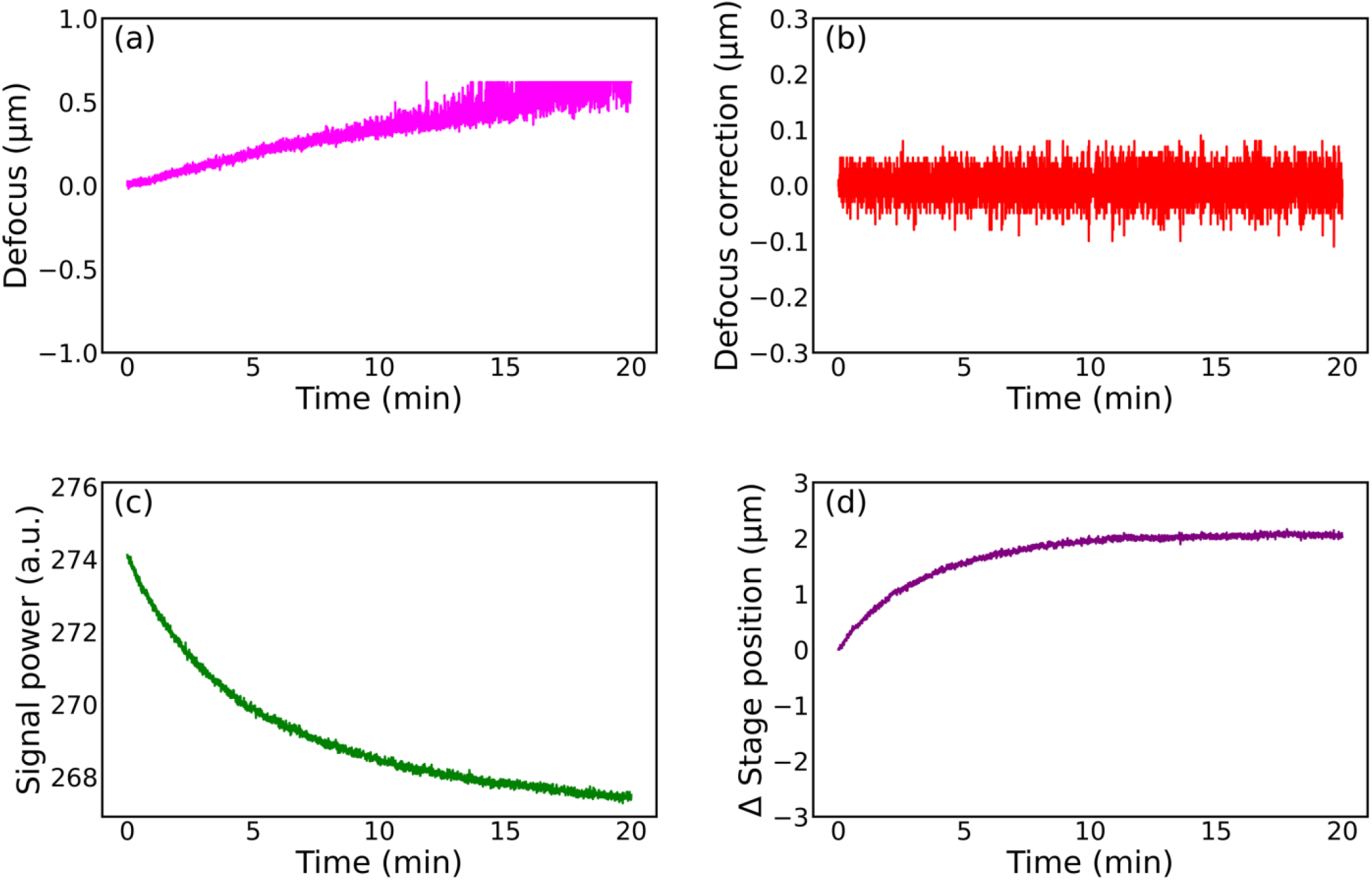
Time-lapse plots during operation of the LED/MMOF openAF autofocus module switched on from cold, revealing a correlation between LED power and focus drift apparent only by astigmatic imaging. (a) the defocus (µm) measured by astigmatic nanobead imaging (magenta), showing >500 nm drift before exceeding the astigmatic imaging range after 20minutes; (b) the required defocus correction (µm) reported by the LED/MMOF openAF module (red); (c) variation in LED power (a.u.) represented by the average intensity of the autofocus camera image (green), showing a ∼1.5 % decrease over the 20-mins warm-up period; and (d) the position (µm) of the piezo-electric z-stage reported by its encoder (purple).

We reasoned that there was a systematic power-dependent error in the calculation of the *openAF* defocus metric that was due to the background signal changing with LED power – noting that the subtraction of the pre-recorded background image was the only intensity-dependent component of the *openAF* calculation of defocus. We therefore compared each real-time autofocus camera image with a reference in-focus image in the autofocus calibration z-stack and used the average intensities to continuously scale the pre-recorded background intensity image before subtracting it from the (real-time) autofocus camera image being used to compute defocus. This effectively normalises the background subtraction to variations in LED power.

As is evident in Figure 5, this normalisation of the background image efficiently eliminated the LED power-dependent error, providing robust operation of the autofocus immediately after the LED was switched on, despite the LED power changing by 2% during the first 30 minutes. In this mode the autofocus accurately maintained focus with a standard deviation of ∼7 nm over 45 minutes. At this point the signal to the z-stage was switched off such that the astigmatic defocus measurement -and the *openAF* defocus measurement - report the focal drift of the uncorrected microscope. This performance will ultimately be limited by the precision of the piezo z-stage (PI, Q-545.140) with controller (PI, E-873.1AT), which has a specified minimum displacement of 6 nm and an encoder resolution of 1 nm, as well as by the precision of the astigmatic imaging measurement.

**Figure 5.**
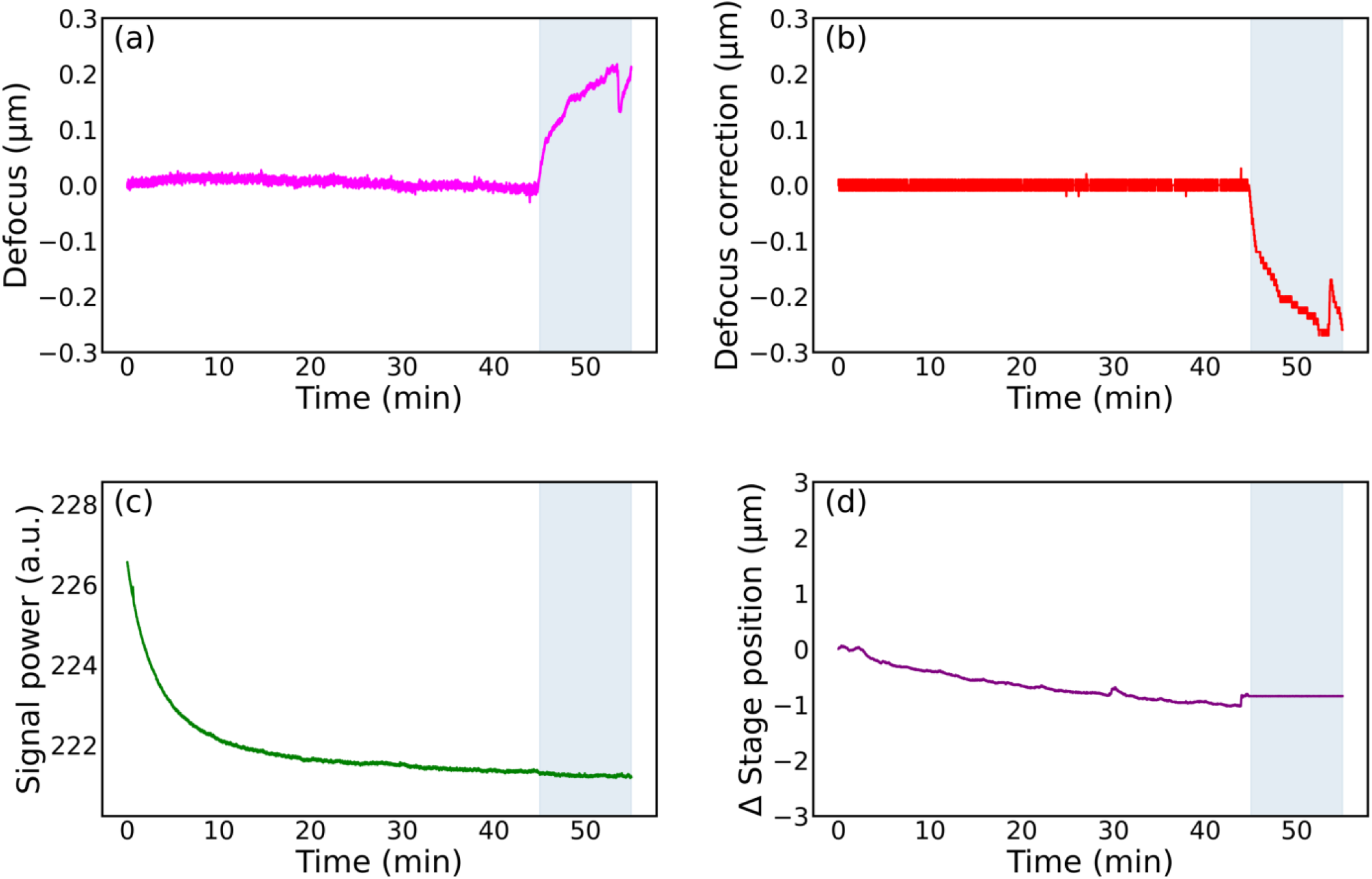
Time-lapse plots during the operation of the LED/MMOF openAF autofocus module switched on from cold with continuous background intensity normalisation : (a) defocus (µm) measured with the astigmatic nanobead imaging (magenta), showing SD of 7 nm over 45 minutes; (b) the required defocus correction (µm) reported by the LED/MMOF openAF module (red); (c) variation in LED power (a.u.) as represented by the average intensity at the autofocus camera (green), showing ∼2% decrease over the 45 min period; and (d) position (µm) of the piezo-electric z-stage reported by its encoder (purple). Note that the z-stage was disabled at 45 minutes.

We applied the same astigmatic fluorescent bead imaging approach to monitor the precision of the *openAF* autofocus module configured with the SLD/SMOF light source. Figure 6 shows how the SLD power at the autofocus camera varied over time, with a ∼5% decrease in power leading to a defocus drift >100 nm –which is lower than for the (unnormalized) LED-based OAF. However, the performance was still improved with the implementation of background intensity normalisation before subtraction, as illustrated in Figure 7, where the focus was maintained with a standard deviation of ∼8 nm over 45 minutes, until the signal to the z-stage was switched off.

**Figure 6.**
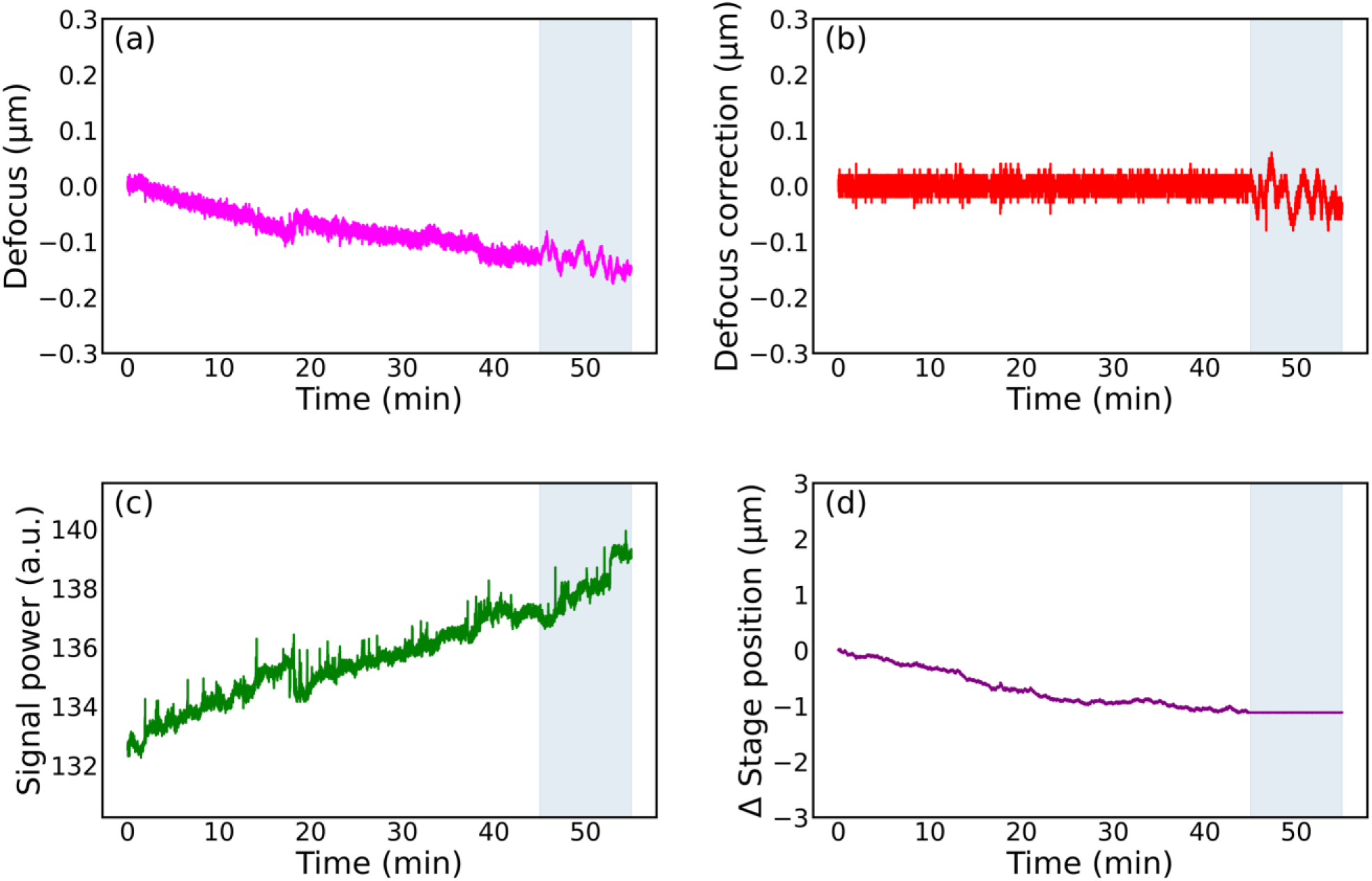
Time-lapse plots of the operation of the SLD/SMOF openAF autofocus module switched on from cold without continuous background intensity normalisation: (a) the defocus (µm) measured with the astigmatic nanobead imaging (magenta), showing drift of >100nm over 45 minutes; (b) the required defocus correction (µm) reported by the SLD/SMOF openAF module (red); (c) the variation in power (a.u.) of the SLD as represented by the average intensity of the autofocus camera image (green), showing an increase of ∼5% over the 45 minutes; and (d) the position (µm) of the piezo-electric z-stage reported by its encoder (purple). Note that the z-stage was disabled at 45 minutes.

**Figure 7.**
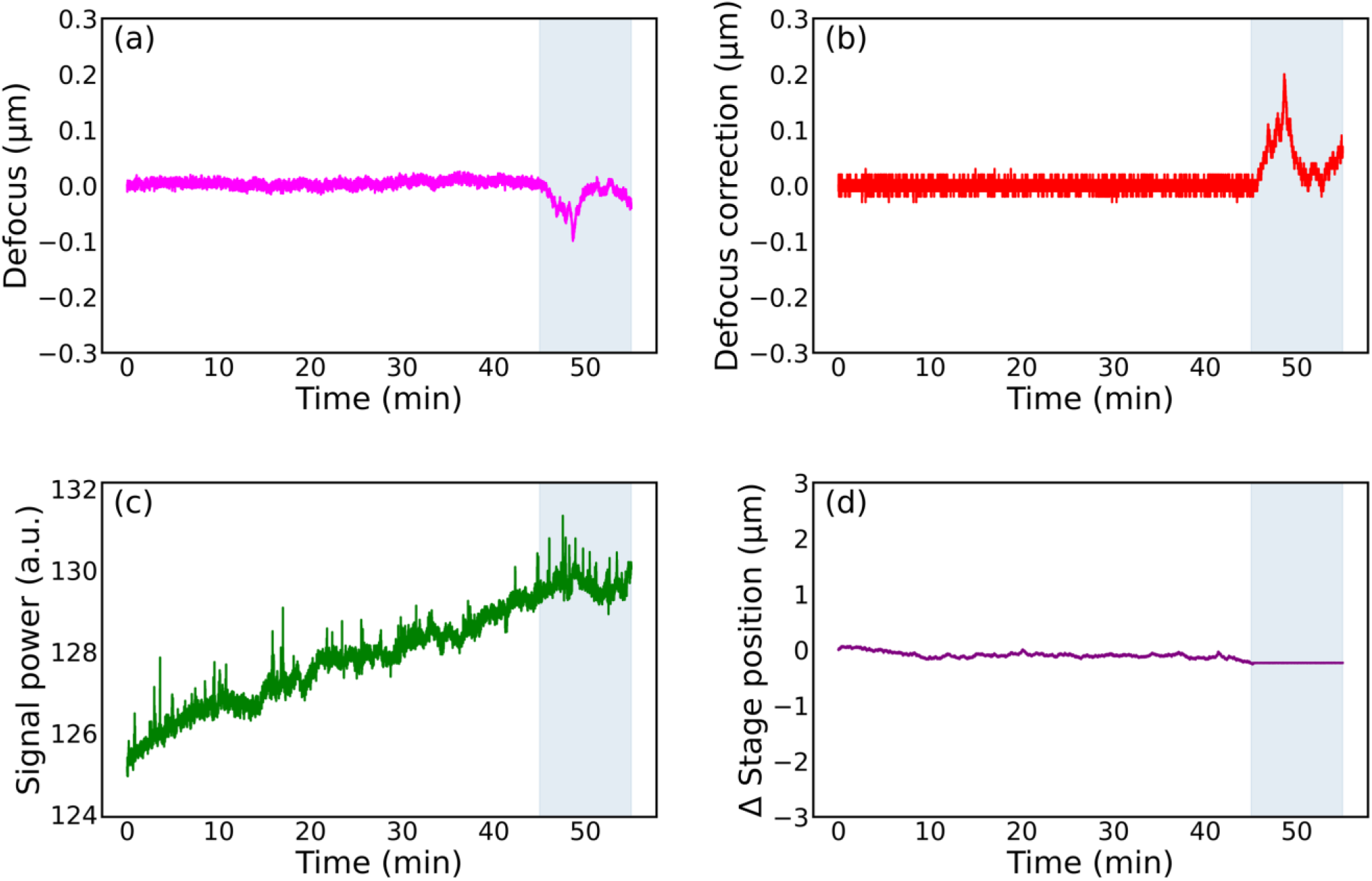
Time-lapse plots of the operation of the SLD/SMOF openAF autofocus module switched on from cold with continuous background intensity normalisation: (a) the defocus (µm) measured with the astigmatic nanobead imaging (magenta), showing SD of 8 nm (b) the defocus correction (µm) reported by the SLD/SMOF openAF module (red), (c) the variation in SLD power (a.u.) as represented by the average intensity at the autofocus camera image (green) and (d) the position (µm) of the piezo-electric z-stage reported by its encoder (purple). Note that the z-stage was disabled at 45 minutes.

To quantify the operating range of the LED/MMOF openAF autofocus system, we implemented a program to translate the objective lens to predetermined values of defocus, with 10 repeated acquisitions at each value. Figure 8(a) shows a plot of the programmed (“true”) defocus against the average defocus value calculated by the *openAF* autofocus module. This indicated the system’s operational range to be 15 µm, with the range being asymmetric about the focus position due to the focal planes of the autofocus camera image and the fluorescence camera image being offset. Figure 8(b) shows a box plot presenting the median and interquartile range of the defocus error for each programmed objective lens displacement. It is apparent that this uncertainty in the predicted defocus is significantly less than the DOF of the 100x 1.4 NA Oil objective lens (UPLSAPO) (200 nm for a wavelength of 550 nm).

**Figure 8.**
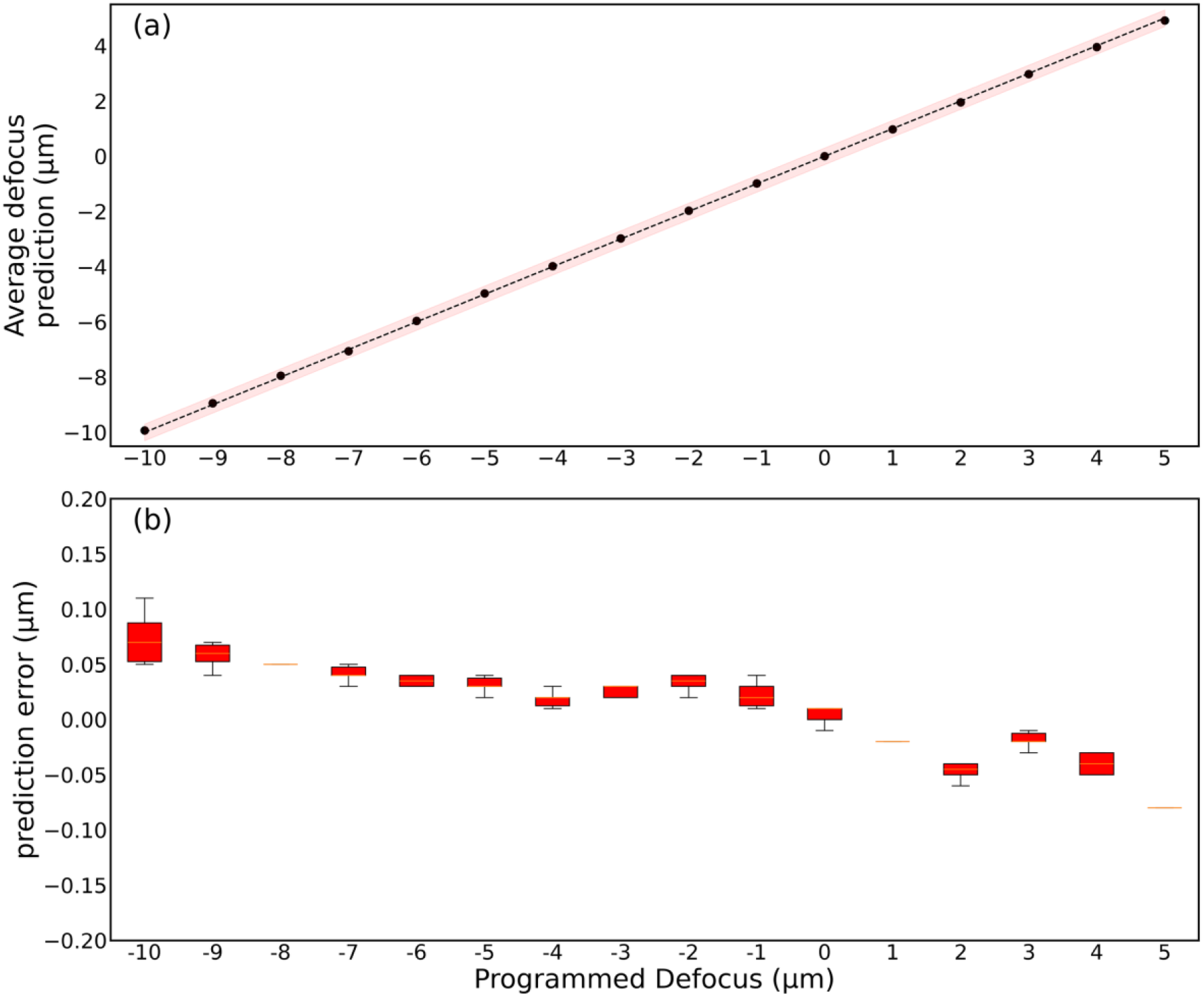
Autofocus range measurement: (a) the average defocus prediction (µm) versus programmed (“true”) defocus for a 15 µm range from -10um to +5um showing that the average prediction remains within the depth of field across the full range; (b) boxplot of prediction errors (µm) versus programmed (“true”) defocus.

The LED/MMOF *openAF* operating range is limited by the rapid expansion of the LED radiation emerging from the MMOF with defocus, which overfills the autofocus camera sensor. When defocus correction required exceeds 15 µm range, which may occur for applications such as multiwell plate imaging, the *openAF* system could employ a two-step correction method, as previously reported^26^, utilising the variation of average intensity at the autofocus camera over the range of the calibration data for the first step defocus metric.

## Conclusions

In conclusion we have presented a straightforward and robust means to quantify the performance of autofocus systems that is based on astigmatic imaging of fluorescent beads analysed using a 2D autocorrelation-based algorithm. We demonstrated this using the open-source *openAF* autofocus module implemented on an *openFrame*-based microscope with a 100x 1.4 NA oil immersion objective lens. We quantify the performance of a new LED/MMOF implementation that offers convenience at significantly lower cost, and without raising laser safety concerns. We have also quantified the performance of the previously published SLD/SMOF configuration.

The accuracy of the astigmatic imaging method to quantify defocus revealed that small variations in the power of the autofocus light source could degrade the performance of the autofocus module. We showed that this could be mitigated by continuously normalising the average intensity of the calibration background image before subtraction from the real time autofocus camera images, enabling the new LED/MMOF implementation to provide SD of <10 nm axial stability over at least 45 minutes. We note that variations in the average intensity at the autofocus camera could arise from changes in the coupling efficiency of light coupled into the SMOF/MMOF, as well as in the emitted power from the LED/SLD.

We believe that this improved LED/MMOF-based *openAF* autofocus module will be useful for SMLM and time-lapse microscopy, noting that it performs well when imaging biological samples with a high-NA oil immersion objective lens that provide the lowest reflectivity (∼ 0.4%) from the coverslip/aqueous medium interface of OAF systems.

The LED/MMOF *openAF* closed loop operating range may be extended beyond 15 µm with further optimisation of the system, e.g., by changing the collimation of the MMOF – noting, however, that using an MMOF with a smaller core diameter would result in less power available at the autofocus camera and therefore a longer autofocus response time, and/or by reducing the focal length of the autofocus tube lens. Alternatively, the calibration data range can be extended, and the average autofocus camera intensity can be used to provide the first step in a 2-step autofocus method. These parameters will be explored in future work.

We have also presented a simple “infinity alignment tool” that is useful for rapidly aligning conventional and astigmatic imaging paths in light microscopes.

## Acknowledgements

The authors gratefully acknowledge funding from the Cancer Research UK ICR/Imperial Convergence Science Centre and Research England GCRF Institutional Award as well as the Imperial College London Impact Acceleration Accounts supported by the Biotechnology and Biological Sciences Research Council (BBSRC EP/R511547/1) and the Engineering and Physical Sciences Research Council (EPSRC EP/R511547/1). This project has also benefitted from support from Cancer Research UK (A28450, A29368), from the Chan Zuckerberg Initiative DAF, an advised fund of the Silicon Valley Community Foundation (grants 2021-234618 and 2023-321240. SH acknowledges a PhD studentship from EPSRC. For the purpose of open access, the authors have applied a CC BY public copyright licence to this submitted manuscript.

The modified *openAF* software is available at https://github.com/imperial-photonics/openAF

The autocorrelation analysis software is available at https://github.com/sara98habte/autofocus-evaluation

The data underlying the figures in this paper is available at https://doi.org/10.5281/zenodo.19349029

