## Supplementary Information for "Quantitative evaluation of LED based optical autofocus module"

Variation of autofocus camera images with defocus

Supplementary video 1 – LED OAF autofocus camera z-stack

Supplementary video 1 shows how the LED/MMOF-based OAF autofocus camera image varies as the microscope objective lens is translated axially from -15 µm to 15 µm. Supplementary Figure 1 shows frames from this z-stack for defocus values of -10, 0 and +10 µm. For comparison, Supplementary Figure 1(b) shows analogous autofocus camera images when the SLD/SMOF are used. Although the images look rather different, the variation with axial translation of the objective lens is reproducible and the same autofocus image metrics can be compared to a calibration data set to determine the defocus.

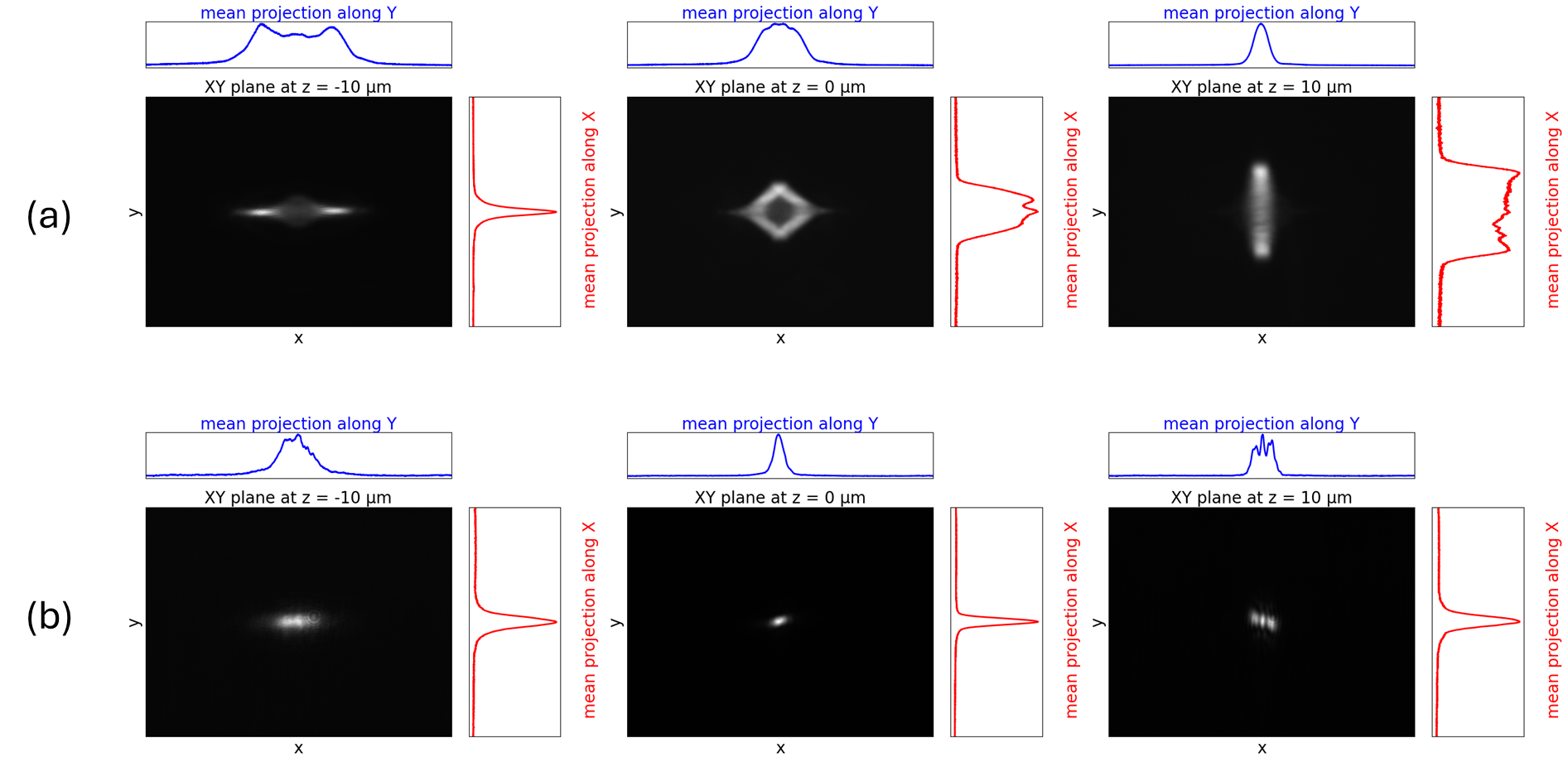

*Supplementary Figure 1. (a) shows LED/MMOF-based OAF autofocus camera images acquired when the microscope objective lens is translated to -10, 0 and +10 µm from the position of best focus(b) shows the SLD/SMOF-based OAF autofocus camera images acquired at the same values of defocus.*

2D autocorrelation of fluorescent bead images to provide independent quantification of defocus.

Supplementary video 2 shows a z-stack from -3 µm to 3 µm of images of TetraSpeck fluorescent beads acquired using the main fluorescence camera of the microscope. Supplementary video 3 shows the corresponding z-stack of the 2D autocorrelation of these fluorescence camera images.

Supplementary video 2 – beads z-stack

Supplementary video 3 – 2D autocorrelation z-stack

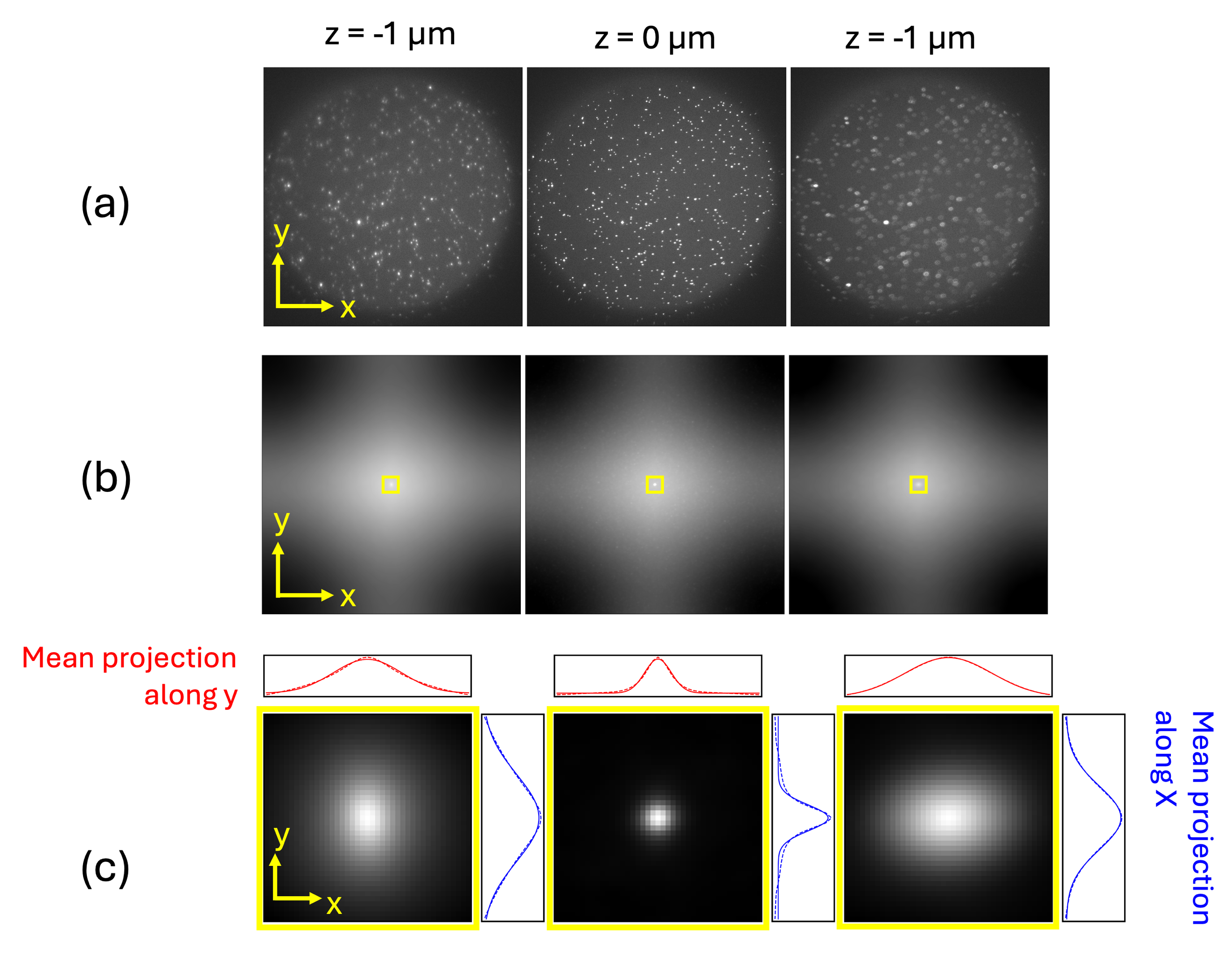

Supplementary Figure 2. (a) fluorescence images of TetraSpeck fluorescent 100 nm beads imaged with 100x oil immersion objective lens acquired at defocus values of -1, 0, +1 µm; (b): respective 2D autocorrelation of these fluorescence images and (c) zoomed in images of central peak of respective 2D autocorrelations functions with plots of mean intensity projections along x and y axes

Supplementary Figure 2(a) shows the fluorescence images of the TetraSpeck fluorescent beads imaged at best focus and ± 1 µm defocus. Supplementary Figure 2(b) shows the 2D autocorrelation function of these images and Supplementary Figure 2(c) show the central peak of the 2D autocorrelation function of the autofocus camera image varies through focus. The 2D autocorrelation function is calculated through the inverse power spectral density of each image given by

$$\begin{aligned} \mathrm{ACF}\left[ m,n \right]= IFFT2D\left\{ \left| FFT2D\left\{ f\left[ m,n \right] \right\} \right|^{2} \right\}\#\left( 1 \right) \end{aligned}$$

Where ACF[m, n] is the discrete autocorrelation at pixel displacement (m, n), f$[m,n]$ is the acquired image, and FFT2D and IFFT2D denote the 2D fast Fourier transform and its inverse.

The defocus of the fluorescence image is determined by fitting the mean projections of the central autocorrelation peak along the x and y axes to a 1D Gaussian model with an offset, given by

$$\begin{aligned} I_{i}=Ae^{\frac{{-\left( i-\mu_{i} \right)}^{2}}{2\sigma_{i}^{2}}}+B\#\left( 2 \right) \end{aligned}$$

Where *i* ∈ {x, y}, *µ_i_* is the mean value for projection *i* and σ is the standard deviation. The offset, *B*, of the 1D Gaussian fit is fitted with the minimum value as an initial estimate. The ratio σ_X_/σ_Y_ is used as a measured of defocus. As discussed on the main text, this metric is compared to a reference data set that is acquired using the following protocol:

**Protocol to acquire astigmatic autocorrelation defocus reference data**

1. Acquire z-stack of 2x2 binned images of fluorescent beads over range of >±1um
2. Calculate 2D autocorrelation function of each autofocus camera image frame
3. Compute orthogonal mean intensity projections of 2D autocorrelation function along X and Y for each frame
4. Fit 1D Gaussian model to X and Y projections for each frame
5. Calculate the ratio of sigma X and sigma Y of fitted mean projections for each frame
6. Generate a look-up table of this ratio versus defocus

**“Infinity alignment tool”**

As discussed in main text, this generally applicable tool can be used to adjust the tube lens and camera in the imaging arm of a microscope – here it was used to set up the astigmatic fluorescence imaging and the autofocus module.

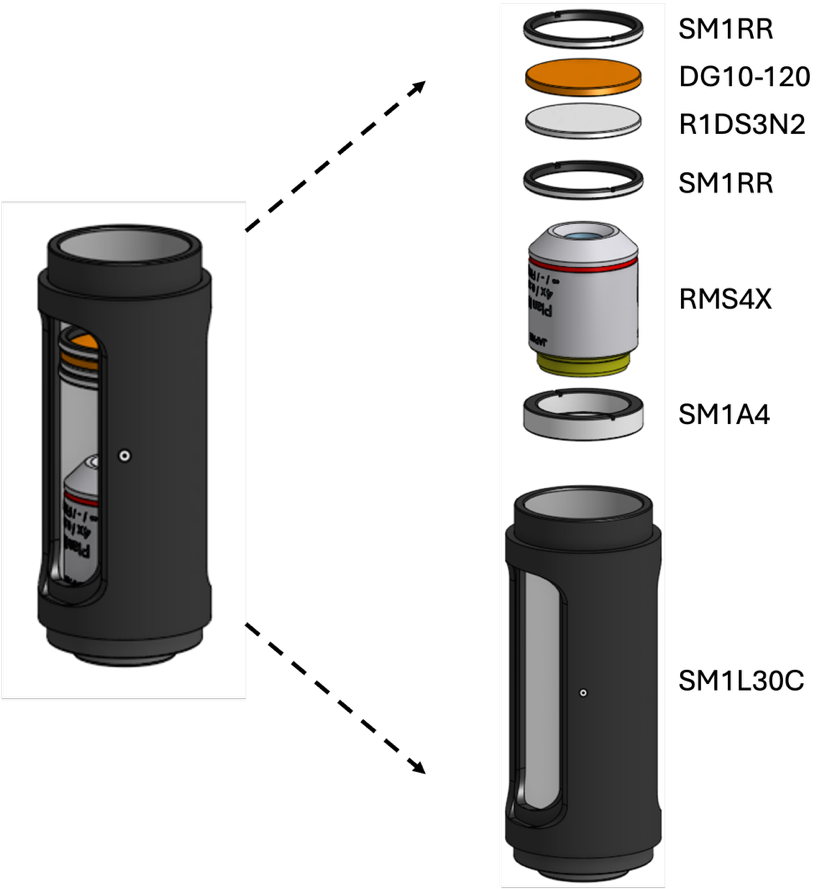

*Supplementary Figure 3. Schematic of “infinity alignment tool” comprising a diffuser, graticule, 25.4 mm diameter aluminium tube with RMS thread adapter and a x4 magnification microscope objective lens. (Thorlabs part numbers are indicated)*

The relative position of the graticule and the microscope objective is critical to provide an image of the graticule at infinity, and this may be precisely set up using an alignment laser (e.g., Thorlabs PL204) and a shearing interferometer (Thorlabs SI050). Subsequently if can be used to quickly set the camera -tube lens distance by removing the main microscope objective and screwing the infinity alignment tool into the objective lens holder. The graticule can then be illuminated from above by any convenient light source and the camera-tube lens distance adjusted until the graticule is in focus.
